# Sensory cues potentiate VTA dopamine mediated reinforcement

**DOI:** 10.1101/2023.10.18.562986

**Authors:** Amy R. Wolff, Benjamin T. Saunders

## Abstract

Sensory cues are critical for shaping decisions and invigorating actions during reward seeking. Dopamine neurons in the ventral tegmental area (VTA) are critical in this process, supporting associative learning in Pavlovian and instrumental settings. Studies of intracranial self stimulation (ICSS) behavior, which show that animals will work hard to receive stimulation of dopamine neurons, support the notion that dopamine transmits a reward or value signal to support learning. Recent studies have begun to question this, however, emphasizing dopamine’s value-free functions, leaving its contribution to behavioral reinforcement somewhat muddled. Here, we investigated the role of sensory stimuli in dopamine-mediated reinforcement, using an optogenetic ICSS paradigm in tyrosine hydroxylase (TH)-cre rats. We find that while VTA dopamine neuron activation in the absence of any external cueing stimulus is sufficient to maintain robust self stimulation, the presence of cues dramatically potentiates ICSS behavior. Our results support a framework where dopamine can have some base value as a reinforcer, but the impact of this signal is modulated heavily by the sensory learning context.

## INTRODUCTION

Environmental cues are critical for successful decision making. Through learning, sensory events associated with rewarding experiences acquire motivational impact, gaining the power to invigorate and reinforce actions on their own. Dopamine neurons in the ventral tegmental area (VTA) are critical in this process, supporting associative learning in both Pavlovian and instrumental settings (Tsai et al., 2009; Witten et al., 2011; Steinberg et al., 2013; Sharpe et al., 2017; Saunders et al., 2018; Keiflin et al., 2019).

Classic studies of intracranial self stimulation behavior demonstrate that animals work hard to receive electrical stimulation delivered to areas of the brain associated with dopamine transmission - including the ventral mid-brain, medial forebrain bundle, and striatum (Olds and Milner, 1954; Mogenson et al., 1979; Corbett and Wise, 1980; Fibiger et al., 1987; Owesson-White et al., 2008). More recently, precise cell-type specific optogenetic targeting has demonstrated that direct activation of dopamine neurons can reinforce self-stimulation behavior (Witten et al., 2011; Kim et al., 2012; Rossi et al., 2013; Ilango et al., 2014; Steinberg et al., 2014). This supports a perspective that dopamine neuron activity can substitute for a natural rewarding stimulus, like food, in the context of learning and reinforcement (Wise, 2004; Berridge, 2007; Steinberg et al., 2013). Recent studies have begun to question the scope of dopamine’s role in this domain, suggesting value-free functions instead, including in novelty, salience, policy learning, and movement kinematics, among other processes (Panigrahi et al., 2015; Coddington and Dudman, 2019; Engelhard et al., 2019; Hughes et al., 2020; Maes et al., 2020; Sharpe et al., 2020; Kutlu et al., 2021; Jeong et al., 2022; Millard et al., 2022; Coddington et al., 2023; Markowitz et al., 2023).

While these various interpretations are not mutually exclusive, it remains somewhat unclear to what extent phasic dopamine neuron activation is itself a direct reward, or if an external sensory element or salient state change is needed for a clear demonstration of ICSS behavior. Determining this is made complicated by the fact that the majority of ICSS studies either include a cue or some other reward paired with stimulation (or fail to report if cues were used), apply ICSS tests after some other learning has occurred, and/or use a targeting method that is not specific to dopamine neurons. Here, we investigated the role of sensory stimuli in dopamine-mediated reinforcement, using an optogenetic ICSS paradigm in naive TH-cre rats (Witten et al., 2011). We found that while brief VTA dopamine neuron activation (1-sec at 20Hz) in the absence of any external cueing stimulus was sufficient to clearly maintain self stimulation, the presence of cues paired with dopamine activation dramatically potentiated ICSS behavior. Our results support a framework where dopamine has some base value as a reinforcer, but the impact of this signal is flexible depending on the sensory learning context. In real world behaving agents, where dopamine neurons are not activated in isolation during learning, dopamine’s role is likely often to assign meaning to associated stimuli and states.

## RESULTS

### Sensory cues accelerate acquisition of dopamine neuron self-stimulation

ChR2-YFP was targeted to dopamine neurons of the VTA in TH-cre+/- rats (**Figure 1, 2A**). Optic fibers were implanted unilaterally above the VTA for optogenetic excitation of dopamine neurons via blue laser light. Fiber placements were clustered over the central VTA for rats in all groups (**Figure 1**).

**Figure 1.**
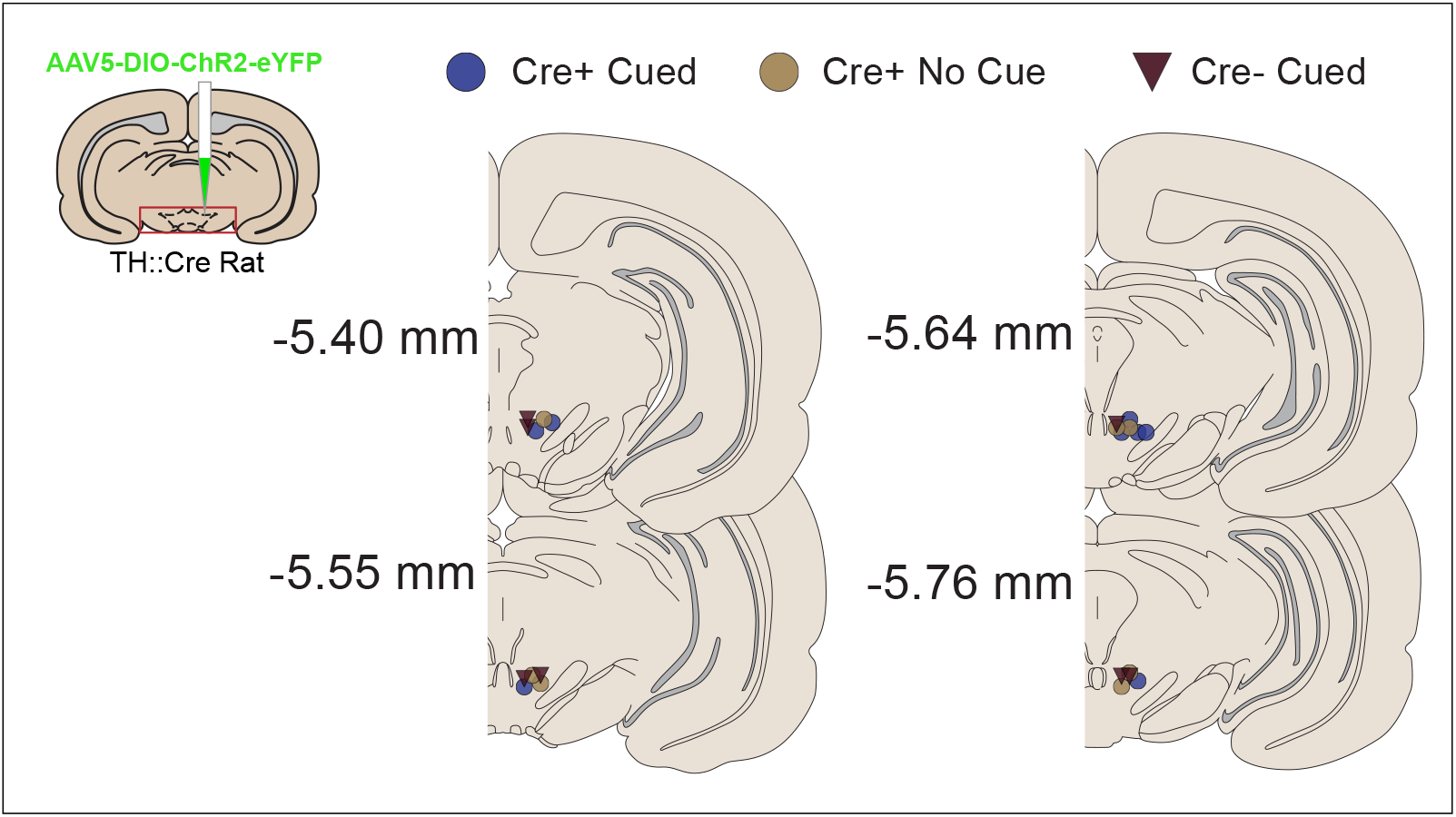
Viral targeting and optic fiber placements in the VTA. ChR2 was targeted to dopamine neurons in the ventral tegmental area dopamine neurons and optic fiber were placed above the VTA for optogenetic manipulations.

**Figure 2.**
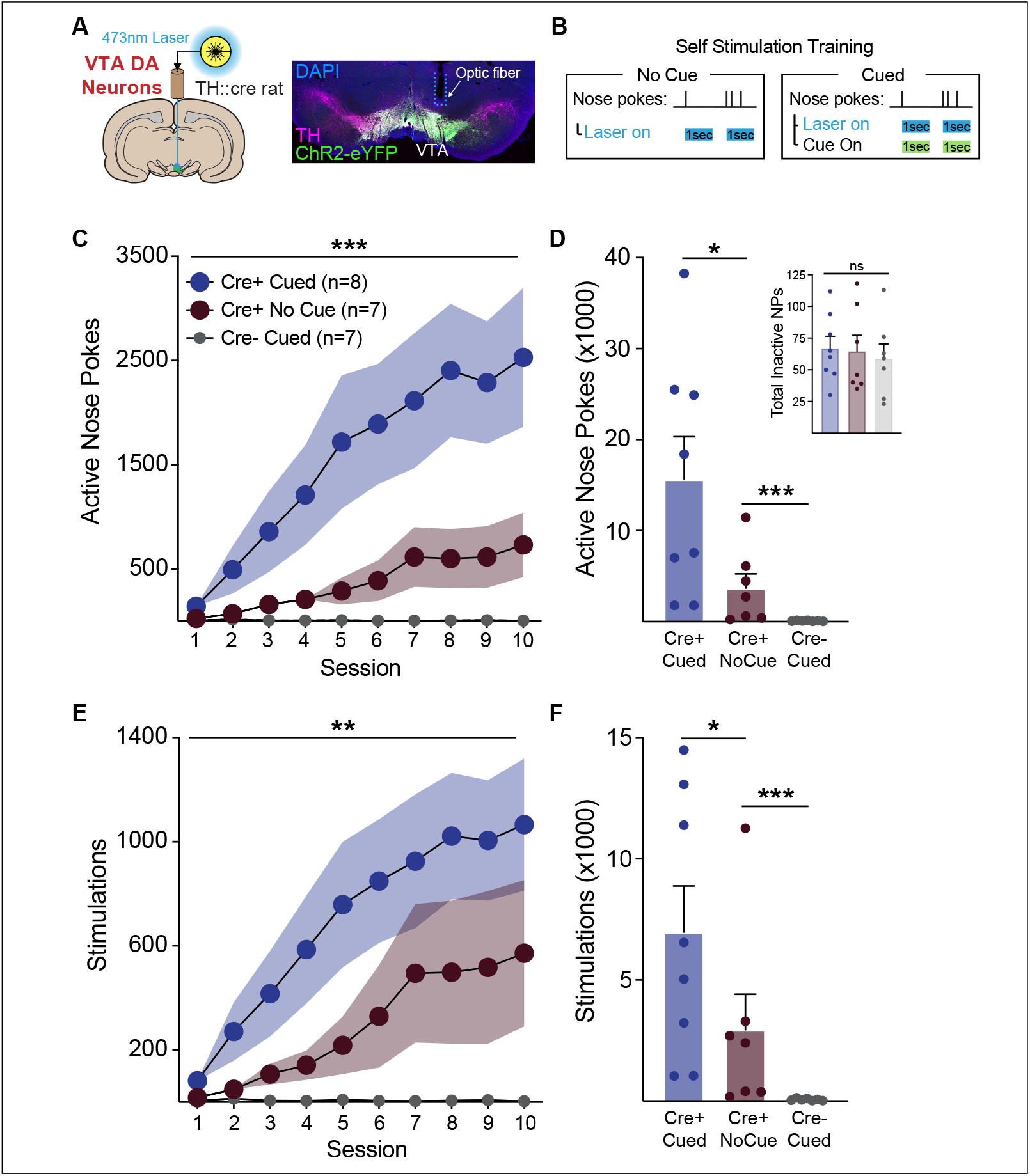
Sensory cues facilitate acquisition of VTA dopamine neuron self stimulation behavior. A) Optic fibers were targeted to the VTA for activation of ChR2-expressing dopamine neurons. B) Rats were allowed to self administer blue laser in 10 1-hour intracranial self stimulation session. Active nose pokes resulted in a 1-sec blue light delivery (20 Hz, 5ms pulses). For rats in the cued groups, this coincided with the illumination of the cue light in the nose poke port, and a delivery of a tone for 1 sec. C) Across training sessions, Cued Cre+ ICSS rats acquired ICSS faster than No Cue Cre+ rats and Cre-controls, as measured by nose poke behavior. D) Cued Cre+ rats completed significantly more nose pokes in total, compared to the other groups. No Cue Cre+ rats completed significantly more nose pokes than Cre-controls. E) Cued Cre+ acquired ICSS faster than No Cue Cre+ rats and Cre-controls, as measured by stimulations received. D) Cued Cre+ rats received more stimulations in total, compared to the other groups. No Cue Cre+ rats received significantly more stimulations than Cre-controls. Responding among the Cre-controls remained low across all sessions. Error represents SEM. ***p<.001, **p<.01, *p<.05.

Animals experienced 10 1-hr intracranial self stimulation sessions, during which they were tethered to an optic fiber connected to a 473-nm laser. Nose pokes into the active port resulted in a 1-sec train of light delivery through the optic fiber (20Hz, 5-ms, 2mW pulses), which was paired with a 1-sec cue (illumination of the active nose port + tone) in the Cre+ Cue and Cre-Cue groups (**Figure 2B**). Cre+ No cue rats received identical laser stimulation, but no cues. Self-stimulation behavior was substantially different across groups (**Figure 2**). The presence of cues accelerated the acquisition of self stimulation, with rats in the Cre+ Cue group making more nose pokes in initial training sessions and reaching greater overall nose poke per session response levels by the end of training (**Figure 2C**; 2-way ANOVA interaction of group and session, F(18,171)=5.92, p<.0001; main effect of group, F(2,19)=7.24, p=.0046). Cre+ Cue rats made significantly more active nose pokes overall, compared to the Cre+ No cue group (**Figure 2D**, Mann Whitney test, p=.020). Cre+ No cue rats responded less for laser stimulation alone (**Figure 2C**), but made significantly more active nose pokes than Cre-Cue control rats (**Figure 2D**, Mann Whitney test, p=.0003) whose responses remained low across all sessions. Across groups, inactive nose pokes were low throughout, and there were no differences in total inactive responses (**Figure 2D** inset, one-way ANOVA no effect of group, F(2,19)=.14, p=.87). Across training, Cre+ Cue (p=.0039) and Cre+ No Cue (p=.0078) rats, but not Cre-Cue (p=.1) rats, discriminated between the active and inactive nose pokes.

We saw a similar pattern for stimulations taken. The Cre+ cued group had the most stimulations across training, finishing with a larger number of stimulations per session, compared to the Cre+ No cue group (**Figure 2E**; 2-way ANOVA interaction of group and session, F(18,171)=3.67, p<.0001; main effect of group F(2,19)=5.92, p=.010). Total stimulations taken were also significantly greater in the Cre+ Cue vs No Cue group (**Figure 2F**, Mann Whitney test, p=.036). Cre+ No cue rats took significantly more stimulations than Cre-cue controls (**Figure 2F**, Mann Whitney test, p=.0003). Together these data indicate that dopamine neuron stimulation can support self-stimulation behavior in the absence of cues. However, self stimulation behavior is substantially accelerated by the presence of cues signaling dopamine neuron activation events.

### Sensory cues potentiate the vigor of dopamine neuron self-stimulation

We next quantified the vigor of self stimulation behavior, by examining the extent to which rats made “extra” nose pokes. During training, in the 1-sec period during which the laser was on for a stimulation event, further nose pokes were counted, but had no additional consequence. Behavior during these “timeout” windows was significantly different across the self stimulation groups (Figure 3). Cre+ Cue rats made many more nose pokes during each stimulation event, as measured by subtracting the number of stimulations from the number of nose pokes across each session (Figure 3A; 2-way ANOVA interaction of group and session, F(18,171)=6.49, p<.0001; main effect of group, F(2,19)=7.09, p=.005). Averaged across sessions, Cre+ Cue rats had significantly greater total nose pokes made during laser delivery timeouts, compared to Cre+ no cue rats (Figure 3B, Mann Whitney test, p=.003). Although they made many fewer than Cre+ Cue rats, Cre+ No Cue rats did significantly more timeout nose pokes compared to Cre-cue controls (Figure 3B, Mann Whitney test, p=.001). We also calculated a ratio score of active nose pokes to stimulations. This revealed that only Cre+ Cue rats consistently showed elevated nose poke behavior above the minimum required to receive stimulations. Across training sessions the nose poke to stimulation ratio increased for Cre+ Cue rats relative to the other groups (**Figure 3C**; 2-way ANOVA interaction of group and session, F(18,171)=2.28, p=.0033; main effect of group, F(2,19)=11.34, p=.0006). Across all sessions, the average nose poke to stimulation ratio was significantly higher in Cre+ Cue versus No cue rats (**Figure 3D**, Mann Whitney test, p=.007). Notably, Cre+ No cue rats were not significantly different from Cre-cue controls (**Figure 3D**, Mann Whitney test, p=.31) in average nose poke to stimulation ratio. These data indicate that the presence of cues paired with dopamine neuron activation potentiate the vigor of self stimulation.

**Figure 3.**
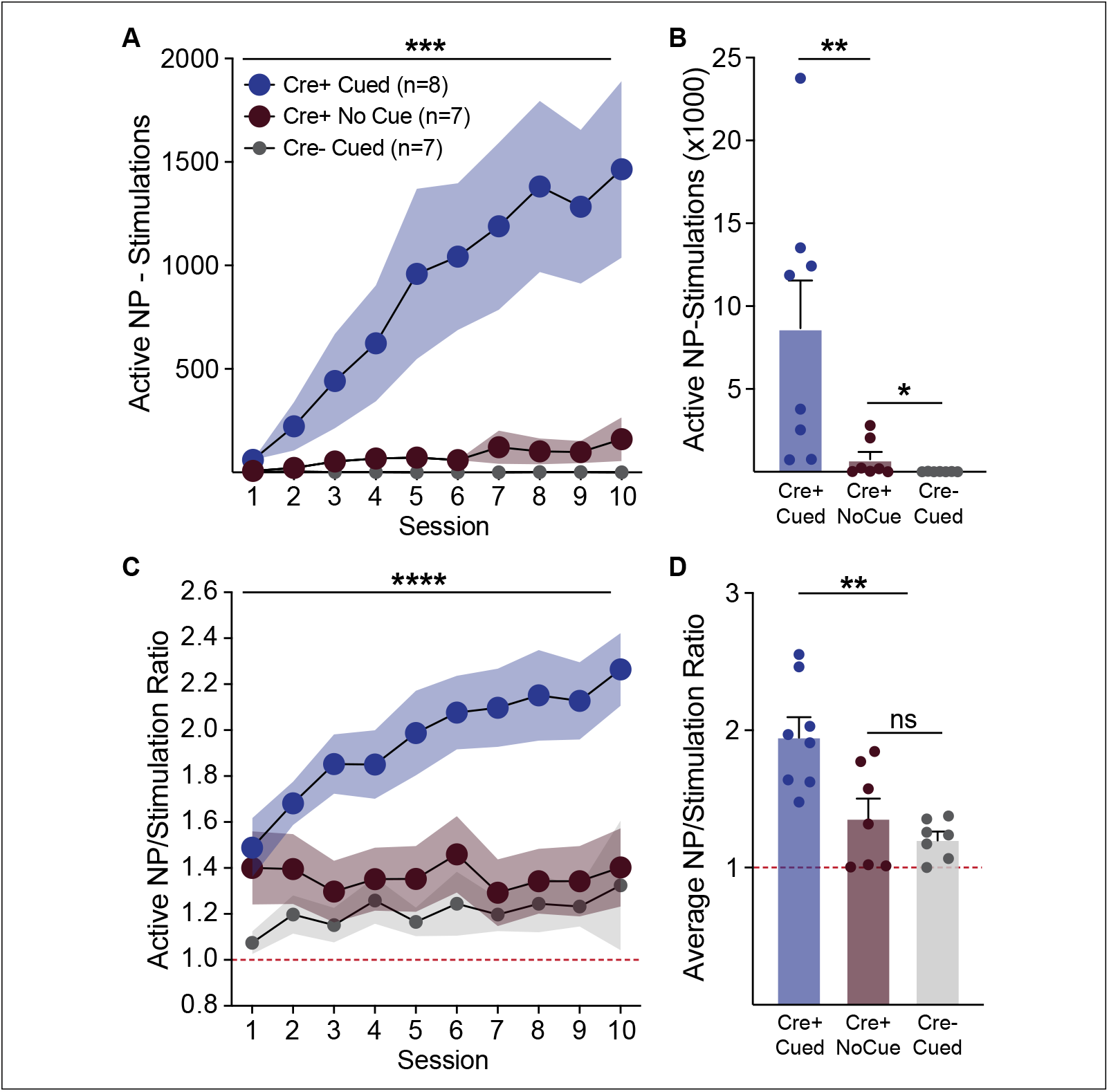
Sensory cues invigorate VTA dopamine neuron self stimulation behavior. Cre+ Cued rats exhibited more vigorous self stimulation behavior, relative to Cre+ No Cued rats and controls. A) This was evident in the elevated number of nose pokes made, compared to the number of stimulations received. Cre+ Cued rats made many more nose pokes during the 1-sec timeout period that coincided with laser delivery, resulting in a positive Active Nose Pokes-Stimulation value across most sessions and B) in total summed across training. In contrast, Cre+ No Cue rats made few extra nose pokes

### Cue history affects rate of ICSS extinction

We next examined the extinction of ICSS behavior (**Figure 4**). Across five sessions, rats were again tethered to an optical cable and allowed to nose poke, where nose pokes had no consequences - no cues or laser were given (**Figure 4A**). Behavior varied substantially across groups under these conditions (**Figure 4**). Rats trained with a cue present during ICSS training maintained a higher level of responding across extinction, compared to the other groups (**Figure 4B**, 2-way ANOVA interaction of group and session, F(8,76)=16.18, p<.0001; main effect of group, F(2,19)=65.85, p<.0001). However, looking at extinction nose pokes as a percentage of the final training session nose pokes, we found that the rate of attrition of behavior in rats with a cue history was more sudden than for rats who did not have a cue during training (**Figure 4C**, 2-way ANOVA interaction of group and session, F(4,52)=2.979, p=.027). This suggests that ICSS behavior in the Cre+ cue group was heavily dependent on the cue’s presence.

**Figure 4.**
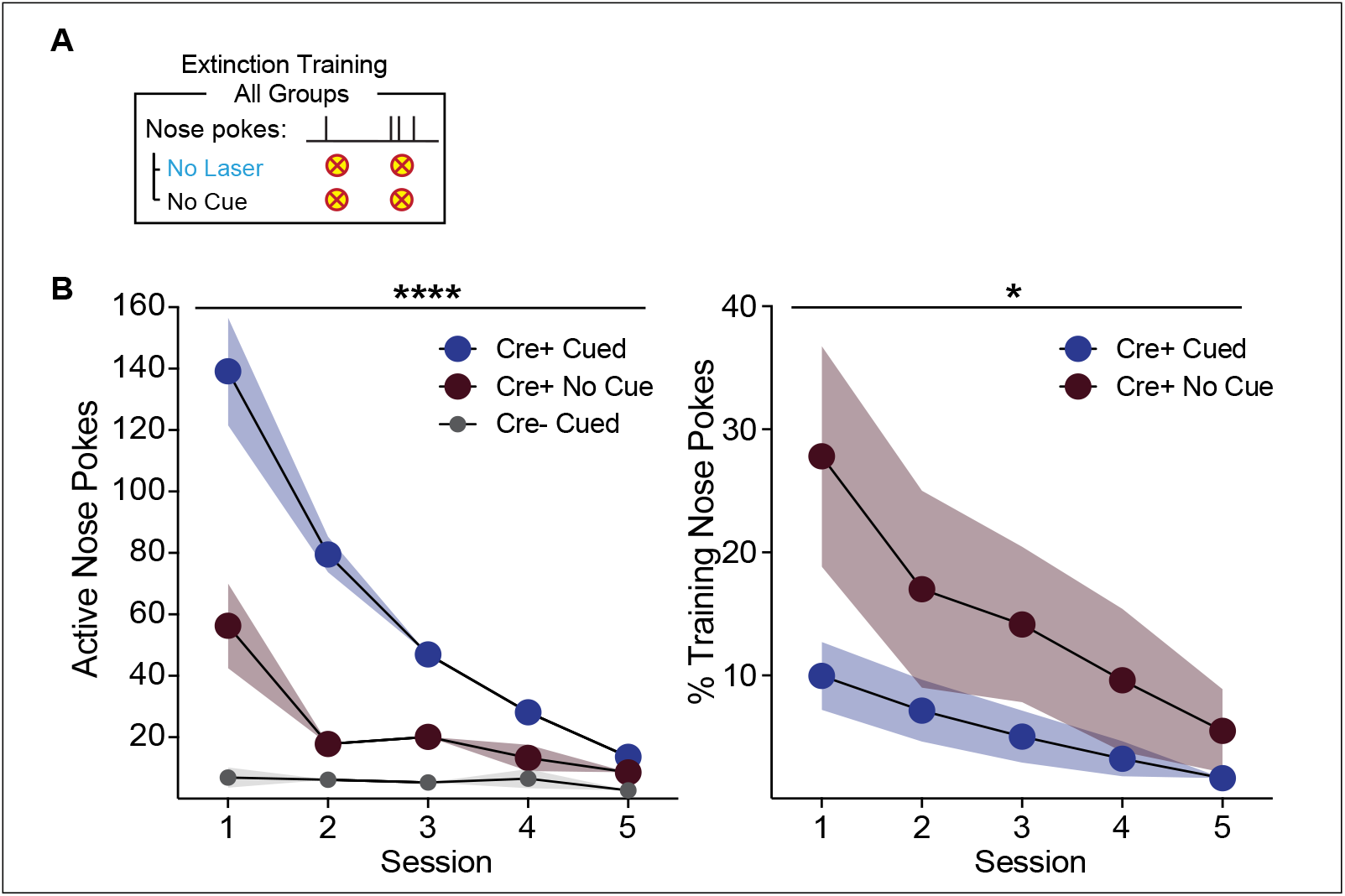
Cue history affects the rate of ICSS extinction. A) Following self stimulation sessions, rats were returned to the testing chambers and allowed to nose poke under extinction conditions, wherein no cues or laser stimulations were delivered. B) Cre+ cued rats maintained higher levels of responding across extinction sessions, relative to Cre+ No cue rats and controls. C) Compared to the final day of ICSS training, extinction of nose poking was more immediate in Cre+ cued rats, relative to Cre+ No cue rats. ****p<.0001, *p<.05.

## DISCUSSION

Here, we demonstrate that dopamine neurons can support ICSS behavior independent of sensory cueing. We found that brief phasic optogenetic activation of VTA dopamine neurons in the absence of any other signaling event was sufficient to produce reliable self-stimulation behavior compared to control animals. Critically, in the presence of cues associated with the receipt of dopamine neuron activation, we observed dramatically potentiated self-stimulation behavior. Rats experiencing dopamine stimulation paired with cues acquired self stimulation more rapidly, took more stimulations, and responded more vigorously than rats receiving optogenetic stimulation alone. Removal of cues under extinction conditions also lead to more rapid attrition of behavior in rats with a history of cue paired during training, relative to no cue rats. Our results add to a long line of research showing that electrical or optogenetic activation of dopamine neurons can support intracranial self stimulation behavior, by clarifying that an unsignaled, brief increase in dopamine activity can act as a value signal akin to a natural reward.

Dopamine neurons in the VTA have long been closely linked to reward learning, motivation, and reinforcement. However, important recent work has begun to question the extent to which dopamine signals reflect behavioral reinforcement as a value signal that substitutes for natural rewards, versus a value-free teaching signal transmitting prediction, salience, novelty, internal state, exploration, or other information (Maes et al., 2020; Sharpe et al., 2020; Kutlu et al., 2021, 2022; Coddington et al., 2023). This work collectively suggests that dopamine itself, divorced from cues, state changes, movement, and other factors, carries no reward-like properties: dopamine serves as a guide of behavior, rather than reinforcing it directly. Our results support a hybrid view. First, we clarify that dopamine neuron activity can directly reinforce actions in the absence of any other sensory or cue-related event, which supports the classic reward value interpretation. We also show strong evidence that this value signal is malleable, and can be potentiated by the presence of sensory cues that have been paired with dopamine neuron activity. This suggests that dopamine neuron signals do not necessarily represent a set static value, but the impact of a given dopamine event is modulated by other features of the learning context, including additional sensory information to support the establishment of associative contingencies. Consistent with this, previous work has shown that in the context of Pavlovian learning, noncontingent, unsignaled dopamine neuron activation does not produce obvious behavioral consequences, including non-specific movement (Coddington and Dudman, 2018; Saunders et al., 2018; Pan et al., 2021). In contrast, if that stimulation event is signaled to the animal with a sensory cue, or if it occurs following a specific action, robust learning and behavioral reinforcement can occur.

Our results suggest that phasic dopamine transmission has the ability to confer some basic value, but that process must interact with other brain processes and states evoked by cues, rewards, and actions for its impact to be fully realized. The extent to which dopamine’s contribution to the net “value” within a learning and decision making context will likely depend on the details - the identity, duration, configuration, and learning history, etc - of those other features. This makes sense as, in real behavioral contexts, where dopamine neurons are never activated in isolation, myriad other processes modulate dopamine neuron activity and function, including sensory, homeostatic, ingestive, and motor control circuits (Redgrave and Gurney, 2006; McCutcheon, 2015; Tian et al., 2016; Coddington and Dudman, 2019; Grove et al., 2022) Thus, value is not solely the domain of dopamine, and in many cases, the value component of dopamine’s contribution to a given behavior state may be negligible (Adamantidis et al., 2011; Maes et al., 2020; Sharpe et al., 2020; Coddington et al., 2023; Markowitz et al., 2023). Collectively, this supports a framework where VTA dopamine serves an important role in credit assignment (what led to that?) and evaluation (how was it?).

Our results are consistent with a recent study (Fraser et al., 2023) demonstrating that VTA dopamine neuron activity supports flexible self-stimulation behavior. Rats will engage in complex seeking tasks, including completing a chain of actions, in order to receive VTA dopamine neuron activation. Further, the presence of cues paired with VTA dopamine neuron activation promotes higher breakpoints on a progressive ratio schedule. This suggests that VTA dopamine neurons can assign value to stimuli and actions, which facilitates both vigorous but also flexible behavior. In contrast, self stimulation of SNC dopamine neurons, while similar to VTA levels for continuously reinforced responding, does not persist when multiple actions are required to produce stimulation (Fraser et al., 2023). Thus while VTA and SNC dopamine neurons may broadly signal a similar base value (Ilango et al., 2014), the learning processes they facilitate are qualitatively different (Wise, 2009; Howe and Dombeck, 2016; Saunders et al., 2018; Keiflin et al., 2019).

In our studies, the presence of a cue dramatically elevated dopamine neuron self stimulation behavior, where some rats made 1000s of responses in one hour. Cued rats not only received many more stimulations than non-cued rats, they nose poked at double the rate. This particular version of self stimulation could be consistent with the development of compulsive-like behaviors thought to underlie addiction and related diseases. Recent studies have shown that individual differences in the extent to which dopamine neuron self stimulation becomes compulsive-like relate to the emergence of unique plasticity within corticostriatal circuits (Pascoli et al., 2015, 2018). Our data suggest that cues play a prominent role in the development of compulsive dopamine seeking. Individual differences in cue responsivity and dopamine activation patterns therefore likely interact to produce different vulnerabilities in these disease states (Flagel and Robinson, 2017).

The conceptual template for VTA dopamine’s functions in learning grows ever more complex, as new tools and models allow for greater precision in manipulation and measurement of the system during behavior. Our results support a view of VTA dopamine as both reward and value-free teaching signal. This work, in combination with several other recent studies, highlights the importance of the learning context in which dopamine’s functions are being probed.

## METHODS

### Subjects

Male and female Th-cre transgenic rats (N=22; 15M, 7F), bred on a Long Evans background, were used in these studies. These rats express Cre recombinase under the control of the tyrosine hydroxylase (TH) promoter (Witten et al., 2011). Wild-type littermates (Th-cre-) were used as controls. After surgery rats were individually housed with ad libitum access to food and water on a 0700 to 1900 light/dark cycle (lights on at 0700). All rats weighed >250 g at the time of surgery and were 5-9 months old at the time of experimentation. Rats were handled by the experimenter several times before and after surgery. Experimental procedures were approved by the Institutional Animal Care and Use Committees at the University of California, San Francisco and at Johns Hopkins University and were carried out in accordance with the guidelines on animal care and use of the National Institutes of Health of the United States. Upon initial investigation, we found no evidence of sex-biased effects, so males and females were collapsed within each ICSS group for all reported data.

### Surgical Procedures

Viral infusions and optic fiber implants were carried out as previously described. Rats were anesthetized with 5% isoflurane and placed in a stereotaxic frame, after which anesthesia was maintained at 1-3%. Rats were administered saline, carprofen anesthetic, and cefazolin antibiotic intraperitoneally. The top of the skull was exposed and holes were made for viral infusion needles, optic fiber implants, and 5 skull screws. A cre-dependent virus coding for ChR2 (AAV5-Ef1α-DIO-ChR2-eYFP, titer 1.5–4e12 particles/mL, University of North Carolina) was infused unilaterally (1 μl at each target site, for a total of 4 μl per rat) at the following coordinates from Bregma for targeting VTA cell bodies: posterior -6.2 and -5.4mm, lateral +0.7, ventral -8.4 and -7.4. Viral injections were made using a microsyringe pump at a rate of 0.1μl/min. Injectors were left in place for 5 min, then raised 200 microns dorsal to the injection site, left in place for another 10 min, then removed slowly. Custom-made optic fiber implants (300-micron glass diameter) were inserted unilaterally just above and between viral injection sites (posterior -5.8mm, lateral +0.7, ventral -7.5). Implants were secured to the skull with dental acrylic applied around skull screws and the base of the ferrule(s) containing the optic fiber. Headcaps were painted black, to block light transmission during laser delivery. At the end of all surgeries, topical anesthetic and antibiotic ointment was applied to the surgical site, rats were removed to a heating pad and monitored until they were ambulatory. Rats were monitored daily for one week following surgery. Optogenetic manipulations commenced at least 4 weeks after surgery.

### Optogenetic Stimulation

We used 473-nm lasers (OptoEngine), adjusted to produce output during individual 5-ms light pulses during experiments of ∼2 mW/mm2 at the tip of the intracranial fiber. Light power was measured before and after every behavioral session to ensure that all equipment was functioning properly. For all optogenetic studies, optic tethers connecting rats to the rotary joint were sheathed in a lightweight armored jacket to prevent cable breakage and block visible light transmission.

### Intracranial Self Stimulation Training

Following recovery from surgery and at least 4 weeks for viral expression, rats were brought to the testing chambers (Med Associates) and connected to an optic cable tether to receive optogenetic stimulation. For all ICSS sessions for all groups, chamber lights were constantly illuminated, to further occlude unwanted light pollution from the laser delivery events. Two nose poke ports were positioned on one side of the chamber. For rats in the cre+ cued and crecued groups, pokes in the active port resulted in a 1-s laser train (20 Hz, 20 5-ms pulses, fixed-ratio 1 schedule with a 1-s timeout during each train), that coincided with the delivery of a cue complex – lights in the nose poke illuminated and a tone played. For rats in the cre+ no-cue group, active nose pokes produced the same 1-s laser train, but no cues. Inactive nose pokes were recorded, but had no consequences.

### Extinction

The day after the final ICSS training session, rats were returned to the testing chambers for the first day of extinction. During this 1-hr session, rats were tethered and nose pokes were measured as before, but laser was not delivered. For rats in the cued groups, nose pokes continued to produce the cues. For rats in the no-cue group, no cue was presented following active nose pokes, as before. Extinction continued for five daily sessions.

### Histology

Rats were deeply anesthetized with sodium pentobarbital and transcardially perfused with cold phosphate buffered saline followed by 4% paraformaldehyde. Brains were removed and post-fixed in 4% paraformaldehyde for ∼24 hours, then cryoprotected in a 25% sucrose solution for at least 48 hours. Sections were cut at 50 microns on a cryostat (Leica Microsystems). To confirm viral expression and optic fiber placements, brain sections containing the midbrain were mounted on microscope slides and coverslipped with Vectashield containing DAPI counterstain. Fluorescence from ChR2-eYFP as well as optic fiber location was then visualized. Tissue from wild type animals was examined for lack of viral expression and optic fiber placements. To verify localization of viral expression in dopamine neurons we performed immunohistochemistry for tyrosine hydroxylase and GFP. Sections were washed in PBS and incubated with bovine serum albumin (BSA) and Triton X-100 (each 0.2%) for 20 min. 10% normal donkey serum (NDS) was added for a 30-min incubation, before primary antibody incubation (mouse anti-GFP, 1:1500, Invitrogen; rabbit anti-TH, 1:500, Fisher Scientific) overnight at 4°C in PBS with BSA and Triton X-100 (each 0.2%). Sections were then washed and incubated with 2% NDS in PBS for 10 minutes and secondary antibodies were added (1:200 Alexa Fluor 488 donkey anti-mouse, 594 donkey anti-rabbit or 647 chicken anti-rabbit) for 2 hours at room temperature. Sections were washed 2 times in PBS and mounted with Vectashield containing DAPI. Brain sections were imaged with a Zeiss Axio 2 microscope.

### Statistics

Behavioral data were recorded with Med-PC software (Med Associates) and analyzed using Prism 9.0. Two-changes in behavior among the groups across training. Mann-Whitney tests were used to compare group averages across sessions. Effect sizes were not predetermined. Rats were included in optogenetic behavioral analyses if optic fiber tips were no more than ∼500 microns dorsal to the VTA. Statistical significance was set at p < 0.05.

